# Population structure reverses selection of variants with proportionally scaled birth and death rates

**DOI:** 10.1101/2025.01.31.636013

**Authors:** Natalia L. Komarova, Dominik Wodarz

## Abstract

A frequently observed phenomenon across the kingdom of life is that a higher reproduction rate can be accompanied by higher mortality. During tumor progression, variants emerge that both reproduce and die faster; faster replicating viruses can be characterized by a faster decay rate; and more frequent pregnancy can be accompanied by a higher chance to die due to predation in ecological systems. Variants with proportionally scaled birth and death rates have been called quasi-neutral mutants. Although life-time reproductive success is not changed, such variants are characterized by fixation probabilities that are somewhat lower (higher) than expected for neutral mutants if birth and death rates are proportionally larger (smaller). Studies were performed in the context of well-mixed populations, and despite the deviation from neutrality, quasi-neutral mutants do not have characteristics of disadvantageous or advantageous mutants, as their fixation probabilities still scale with their initial fractions. Here, we report that in deme-or spatially structured populations, variants with proportionally increased (decreased) birth and death rates become truly disadvantageous (advantageous), and calculate their effective fitness. Furthermore, if mutants have a higher life-time reproductive output than the wild-types and are thus advantageous, a proportional increase of birth and death rates can render them strongly disadvantageous, and vice versa. This changes our understanding of how life-time reproductive success correlates with selection, and has implications for evolutionary dynamics across a range of biological systems.

## Introduction

In many biological scenarios, an increased reproduction rate can be associated with a higher death rate of the organism, especially in microbial and cell populations, due to a cost associated with reproduction. For example, studies have revealed positive correlations between the rate of reproduction and the decay rate of phages that infect the bacterium *Escherichia coli* [1, 2]. Among a variety of tumors, positive correlations between proliferative and apoptotic measures have been reported, and tumor progression appears to be associated with the emergence of cells characterized both by an increased proliferation and death rate [3-9]. In ecological settings, reproduction can increase the chances of death, e.g. through predation [10].

Previous theoretical work has shown that so called quasi-neutral mutants are characterized by non-trivial evolutionary dynamics [11-13]. Quasi-neutral mutants are defined by having proportionally scaled birth and death rates compared to the wild-type, i.e. *r*_*m*_ *=* τ*r*_*w*_ and *d*_*m*_ *=* τ *d*_*w*_, where *r* denotes the reproduction rate, *d* the death rate, subscripts *w* and *m* denote wild-type and mutant individuals, and τ is a constant acceleration or deceleration factor, which we will refer to as a turnover factor. Although the total reproductive output during the life-span of the individuals is identical for wild-type and mutant populations, the mutants are characterized by a fixation probability that is different than that predicted for a neutral mutant [11-13]. If the mutants replicate and die faster than the wild-type population, we refer to them as having a higher turnover than the wild-type population (τ *>1*). In this case, the mutant fixation probability is lower than that of a neutral mutant. The reason is that a faster turnover leads to more demographic fluctuations of the mutant population over a given time interval compared to the wild-type population. This makes the mutant population more prone to extinction, accounting for their lower fixation probability. They, however, do not act like truly disadvantageous mutants because the fixation probability relative to that of a neutral mutant does not decrease with population size. Similarly, decelerated variants (τ <1, a lower turnover than WT) are characterized by a higher fixation probability compared to neutral mutants. This phenomenon was reported for the Verhulst-type (non-constant population) death-birth process [11-13] and for the Moran process [14].

While the assumption of quasi-neutrality is somewhat artificial (because birth and death rates will rarely be changed by exactly the same amount in a biological system), these results indicate that higher turnover of mutant cells can make them less likely to rise and displace a resident population with a slower turnover at steady state. This can be counter-intuitive especially because during population growth at lower densities, the higher turnover mutant population is characterized by a faster rate of expansion (due to its faster reproduction rate) [11].

The studies performed so far were made under the assumptions of well-mixed populations. The majority of population interactions, however, involve spatial structures, which can potentially change the outcome of dynamics. Here, we re-visit the evolutionary dynamics of quasi-neutral mutants in the context of spatially structured populations.

## Results

We first consider a deme model where logistic growth dynamics are simulated stochastically with Gillespie-type simulations in each deme [15], and where division can also result in the offspring individuals migrating to other demes, regardless of their location. In each deme, the population fluctuates around a steady state, see Materials and Methods for the details of this modeling approach. In previously published work assuming well-mixed populations, the fixation probability of a quasi-neutral mutant was calculated (for constant populations [14] and for populations with demographic fluctuations, in the diffusion approximation limit [11]). Remarkably, in both cases, the fixation of a mutant, starting from a single individual in a population of (mean) size *N*, is given by 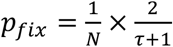, where τ is the fold change in the turnover (τ *>1* for mutant with higher turnover). This fixation probability is lower than that for a neutral mutant (*1/N*) for mutants with higher turnover (τ >1) and it is higher than neutral for mutants with lower turnover (τ <1). The amount of reduction in the fixation probability compared to a neutral mutant, however, does not depend on *N* in this scenario: *N x p*_*fix*_ =2/( τ +1) remains constant. For τ >1, this is in contrast to a truly disadvantageous mutant, for which the value of *N x p*_*fix*_ declines exponentially with population size, *N*. For τ <1, again it is different from a truly advantageous mutant, whose fixation probability tends to a constant in a large-*N* limit (and does not scale with 1/*N*). Hence, the term quasi-neutral.

The deme-structured model, however, displays fundamentally different properties. In this setting, the value of *N x p*_*fix*_ for faster turnover mutants declines exponentially with *N* (Fig 1A), which is the characteristic of truly disadvantageous mutants. In other words, although the reproductive success of the higher turnover mutant individuals during their life-span is identical to that of wild-type individuals, they experience negative selection. The extent of the disadvantage the mutant experiences becomes larger with the amount by which the turnover (i.e. the proportionally scaled birth and death rates) is increased relative to the wild-type (parameter τ, Fig 1B), as well as for larger death-to-birth ratios of the individuals (Fig 1C). Note that the relationship between the mutant fixation probability and the death-to-birth ratio of individuals is fundamentally different for the quasi-neutral non-spatial system (increasing with the death-to-birth ratio, *d*_*w*_*/r*_*w*_) and the deme-structured system (decreasing with the death rate-to-birth ratio, *d*_*w*_*/r*_*w*_, Fig 1C). Similar considerations apply to mutants that are characterized by a lower turnover compared to the wild type (τ <1), see supplementary figure S4. In particular, as the population size increases, the probability of fixation increases and saturates at a constant level, indicating that these types of mutants experience a true selective advantage. Therefore, the deme population structure fundamentally changes the properties of mutants in which birth and death rates are proportionally scaled. While in mixed populations, such mutants are quasi-neutral, in deme structured populations mutants with higher (lower) proportionally scaled birth and death rates experience negative (positive) selection.

**Fig 1.**
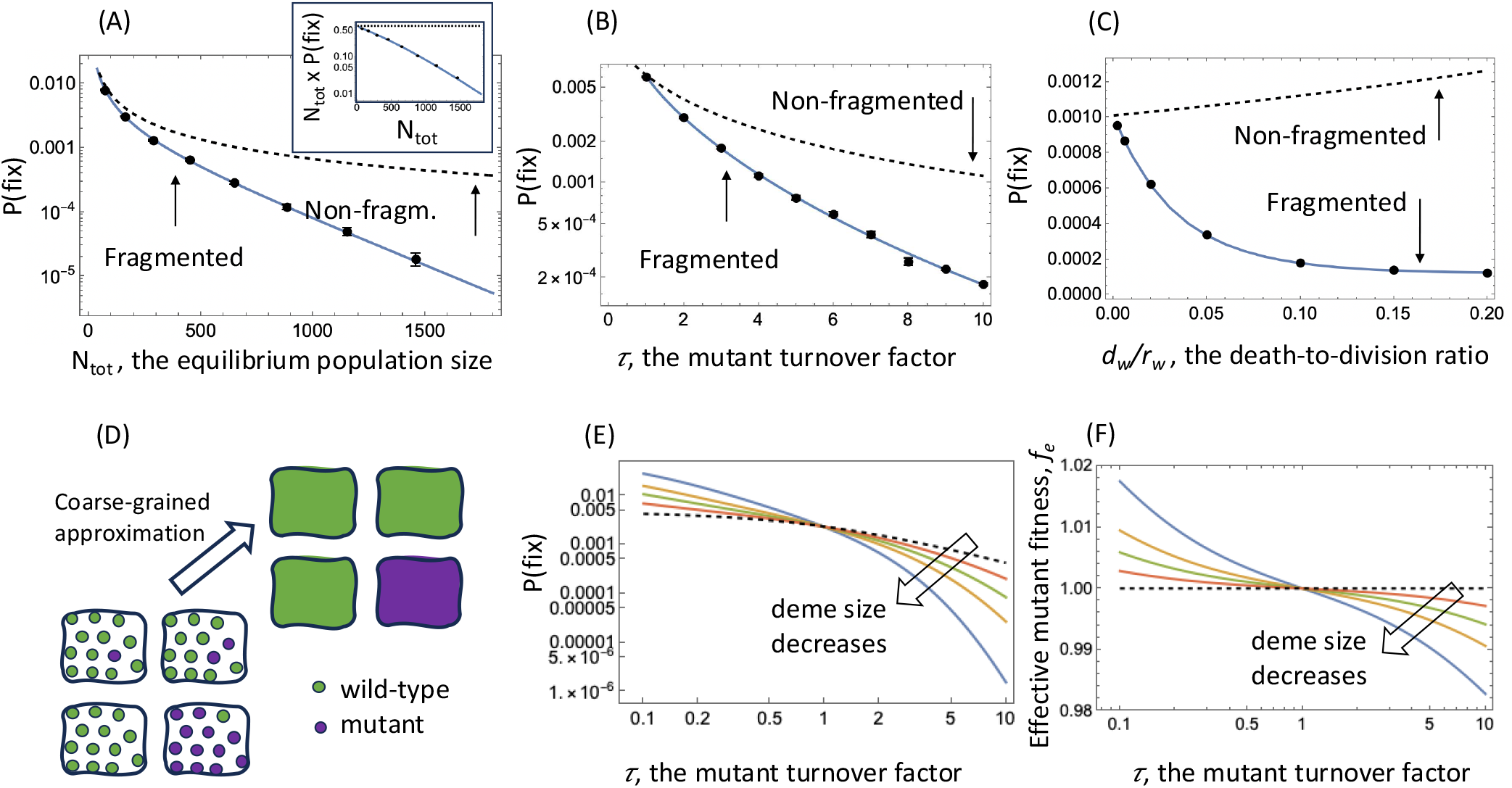
Properties of a mutant with proportionally increased birth and death rates (expressed by turnover factor τ > 1) in the deme-structured model. (A-C): The points show results from computer simulations, including error bars based on more than 10^6^ simulations per point. The solid line is the coarse-grained theory prediction. The dashed line indicates the theory predictions for quasi-neutral mutants in a non-fragmented population. (A) The mutant fixation probability declines exponentially with the total quasi-equilibrium population size, N_tot_, a characteristic of a disadvantageous mutant. Inset: The quantity N_tot_ xP(fix) as a function of N_tot_ is a constant in non-fragmented populations, and declines in fragmented populations. (B) The mutant fixation probability declines substantially with the amount by which the birth and death rates are proportionally increased, τ. (C) The reduction of the mutant fixation probability in the deme-structured compared to the non-spatial population becomes more substantial with higher death to birth ratios. Base parameters were chosen as follows: For (A) *τ=2*; *D* varies from 4 to 100. For (B) *D=9*. For (C) *1=10, D=9*. (D) A schematic showing the concept of a coarse-grained approximation. (E) The probability of mutant fixation (obtained by the coarse-grained approximation) as a function of τ, in a deme-structured model with the total carrying capacity 480, split into equal demes of size K=20,30,40,60. (F) The effective mutant fitnessti f_e_, corresponding to the same deme systems as (E). The rest of the parameters, unless otherwise noted, are *d*_*w*_*/ r*_*w*_*= 0*.*1*; *ε=10*^*-4*^; *K=20*; s=0.

To explain this behavior, we used a coarse-grained approach [16], which is applicable when the mutation rate is sufficiently low such that individual demes are typically homogeneous with respect to the type (either fully wild-type and fully mutant, see the schematic in Fig.1D). In this case, the process is reduced to a 1D Markov walk, governed by a deme conversion process, whereby an individual of one type migrates to a random deme of a different type and fixates successfully in the target deme (see Supplementary Materials, Section 1 for details). Let us denote by *ρ*_*m*_ the probability of a single mutant to fixate in an isolated wild-type deme, and *ρ*_*w*_ (used below) is the probability of a single wild-type individual to fixate in an isolated mutant deme. Then the probability of mutant fixation in a system of *D* demes, starting from a single mutant individual, is given by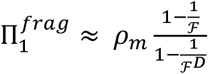, where ℱ is the relative fitness of a mutant deme, defined by the ratio of two deme conversion rates, from wild-type to mutant and from mutant to wild-type. For quasi-neutral mutants,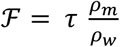. The factor τ accounts for the fold-difference in mutant migration rate (assumed to scale with turnover parameter, τ) and the factor 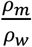 is the fold-difference in fixation rates. Figure 1(A-C) shows excellent agreement between this theory and computer simulations.

Note that the size of the effect depends on the degree of fragmentation and gets stronger as the population is split into a large number of small demes, see Fig 1E and Fig S5. If, on the other hand, the demes are large, such that a diffusion approximation is valid, we have 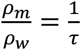, thus resulting in ℱ=1 (deme neutrality) and 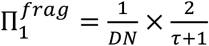, which is exactly the same as the probability of fixation in an non-fragmented system (of size *N*_*tot*_ *=DN*). In other words, fragmentation into large demes does not lead to our reported results. In the case of smaller demes of equilibrium size *N*, we can define effective fitness parameter for quasi-neutral mutants, *f*_*e*_ = ℱ^1/*N*^ (see Fig. 1F). Then, for the probability of mutant fixation in a fragmented population, we have 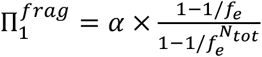, where *α* is a constant independent on the deme number. For faster mutants (τ >1), a deviation from the diffusion limit leads to 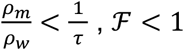, and *f*_1_ < 1, thus making quasi-neutral mutants disadvantageous and resulting an exponential decay of the mutant fixation probability as a function of deme number. For slower mutants (τ <1), we have 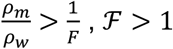, and *f*_1_ > 1, that is, mutants are now advantageous and their fixation probability increases to a constant as a function of *N*_*tot*_. This explains the numerical observations described above.

A biologically more realistic scenario is to consider a mutant that has a higher or lower life-time reproductive output compared to the wild type, but also has proportionally scaled birth and death rates (altered turnover). For example, we can assume that division and death rates of mutants are given by *r*_*m*_ *=* τ *(1+s)r*_*w*_ and *d*_*m*_ *=* τ *d*_*w*_, where subscripts *m* and *w* stand for wild-type and mutant individuals, and where *s* is the selection coefficient. Thus, we have a mutant with proportionally scaled birth and death rates (expressed by the value of τ), but the birth rate is changed further by a factor *1+s*. If *s>0*, the life-time reproductive success of the mutant is larger than that of the wild-type (traditionally considered an advantageous mutant), and for *s<0*, the life-time reproductive success of the mutant is smaller than that of the wild-type (traditionally considered a disadvantageous mutant). We investigated the fixation probability of such mutants.

Figure 2A plots the mutant fixation probability as a function of the fold increase in mutant turnover, τ, for a scenario where *s=0*.*01*. For τ =1 (no change in turnover), the fixation probability is well predicted by the established theory given a 1% fitness advantage (for large populations yielding the probability of fixation simply equal to *s*). As the mutant turnover, τ, is increased relative to the wild-type, however, the fixation probability declines sharply and becomes lower than *1/N*, the fixation probability of a neutral mutant. In other words, by proportionally increasing the birth and death rate of a mutant (turnover parameter τ), the mutant turns from having a selective advantage to having a strong selective disadvantage. Conversely, a mutant that is disadvantageous for τ *=1* (e.g. *s=-0*.*01*) can become advantageous if it has proportionally reduced birth and death rates (τ *<1*, Figure 2B). In Supplementary Materials we provide general expressions for the deme fitness, ℱ, for any type of mutants, under a variety of model assumptions, including different models of migration and mutant initialization. Figure 2(A,B) again demonstrates excellent agreement between the coarse-grained theory and computer simulations in a deme model.

**Fig 2.**
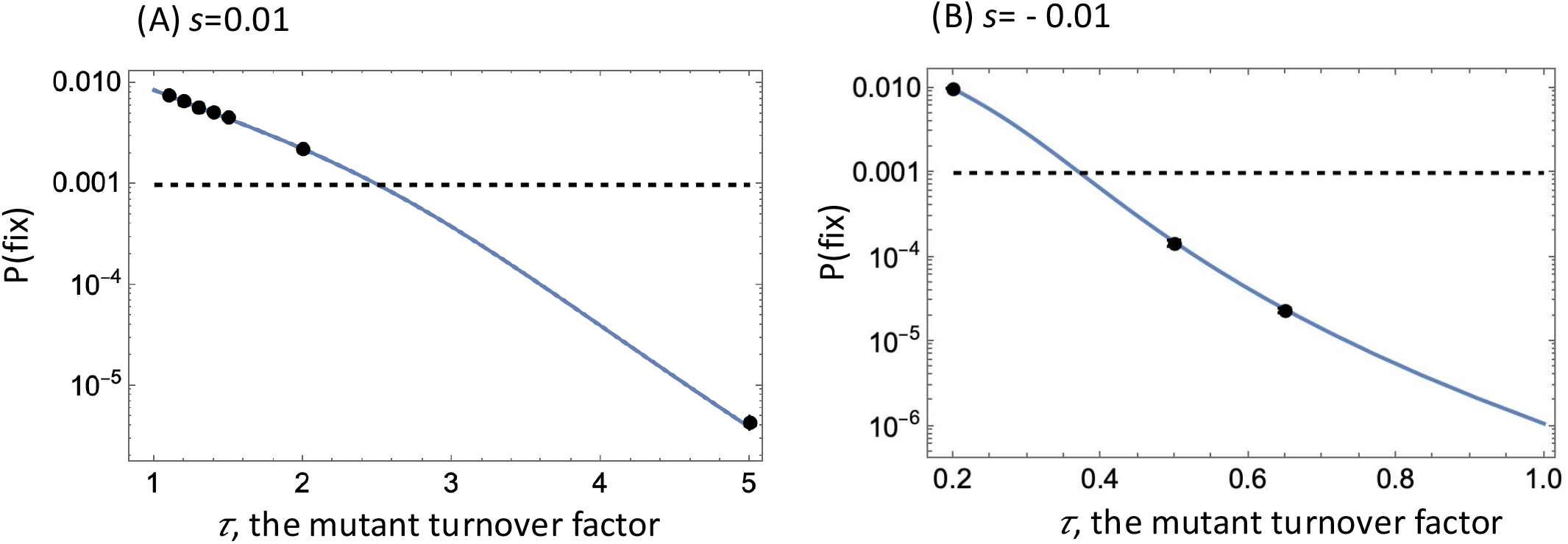
Properties of a mutant with proportionally increased birth and death rates (expressed by turnover factor τ), and additionally a changed life-time reproductive output compared to the wild type in the deme-structured model. The changed life-time reproductive output of the mutant is expressed by the selection coefficient, *s*, such that *r*_*m*_ = τ *(1+s) r*_*w*_. Mutant fixation probabilities are shown as function of τ. The points represent the mean of computer simulations, including error bars based on more than 10^6^ runs. The solid line is the coarse-grained theory prediction. The dashed line represents the fixation probability of a neutral mutant, 1/N_tot._ (A) A mutant that has a 1% higher life-time reproductive output compared to the wild-type is advantageous for τ=1 and small values of τ, but turns disadvantageous for higher mutant turnover rates (larger τ). (B) A mutant that has a 1% lower life-time reproductive output compared to the wild-type is disadvantageous for τ=1 and higher values of τ, but turns advantageous for lower mutant turnover rates (smaller τ). The rest of the parameters are D=64; K=20; *d*_*w*_*/ r*_*w*_*= 0*.*2*; *ε=10*^*-4*^.

The same trends are shown in computer simulations that start with only a wild-type population at equilibrium and assume that wild-type individuals generate mutants at a constant rate μ. Figure 3A shows that a mutant that is advantageous (*s*=0.01) for τ =1 invades the population as expected, but an accelerated mutant with the same relative advantage (τ >1) persists at a selection-mutation balance, a characteristic of a disadvantageous mutant. Figure 3B shows a similar simulation for *s*=-0.01. Without increased turnover (τ =1), the mutant is disadvantageous and persists at a selection-mutation balance. For τ <1, however, the mutant invades the population and fixates, which is characteristic of an advantageous mutant. The coarse-grained approach allows for an analytical estimate of the selection mutation level and the threshold fold-difference in the turnover rate that reverses selection (by making advantageous mutants disadvantageous and the other way around), see Supplementary Materials, Section 2.

**Fig 3.**
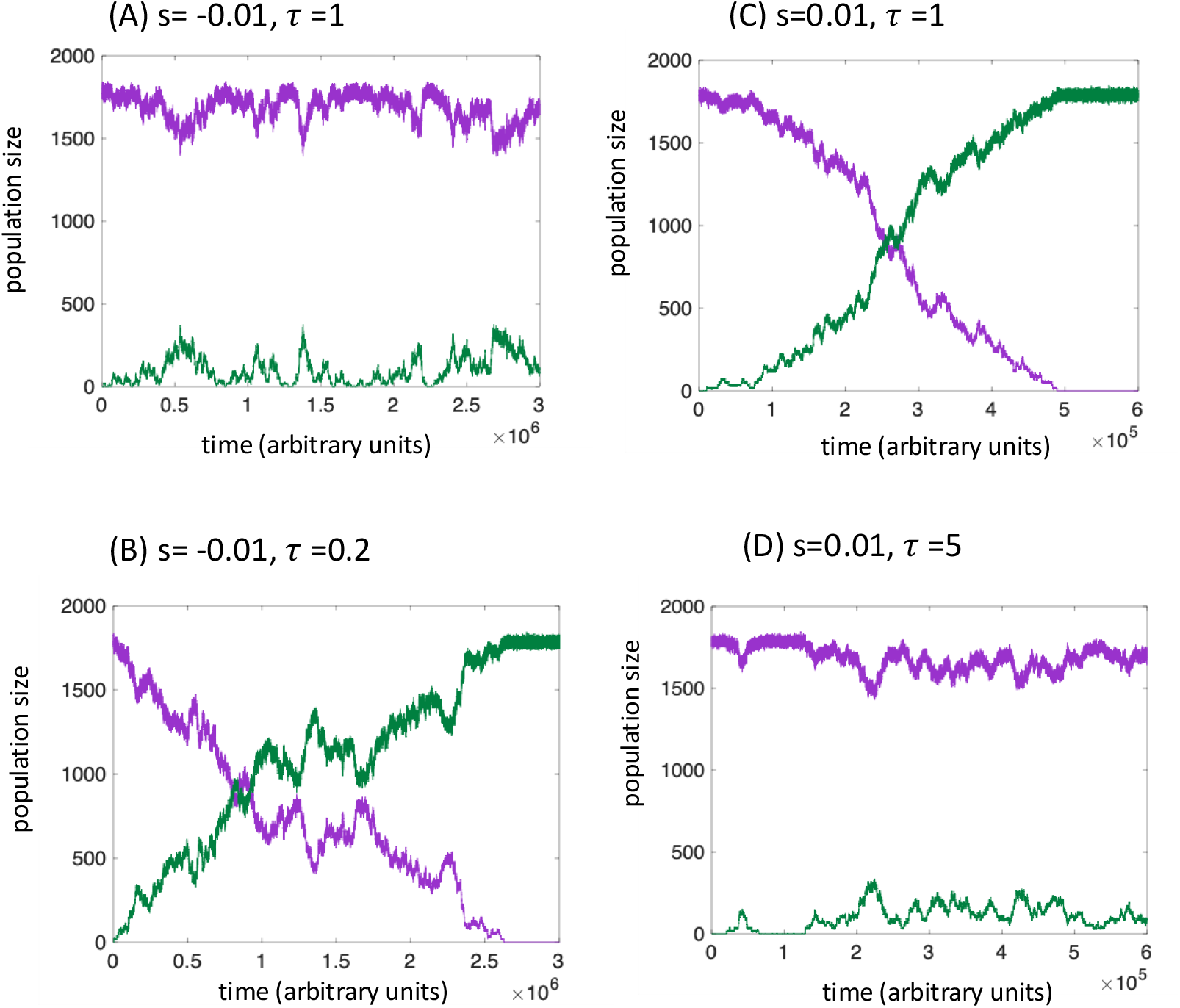
Proportionally scaled birth and death rates can reverse selection in the deme-structured model with mutations. Individual evolutionary trajectories are shown. (A) A mutant with a 1% disadvantage (s=-0.01) is assumedti without any scaled birth and death rates compared to the wild-type (τ=1). As expected, the mutant persists at selection-mutation balance. (B) Same 1% disadvantage, but the mutant simultaneously has 5-fold reduced turnover compared to the wild-type (τ=0.2), leading to a selective advantage. (C) A mutant with a 1% advantage (s=0.01) is assumed, without any scaled birth and death rates compared to the wild-type (τ=1). As expected, the mutant invades. (D) Same 1% advantage, but the mutant simultaneously has 5-fold higher turnover compared to the wild-type (τ=5), and this renders the mutant disadvantageous, persisting at a selection-mutation balance. Baseline parameter values were as follows. *r*_*w*_*=r*_*m*_*=5*; *d*_*w*_*=dm=0*.*5*; *ε=10*^*-4*^; *K=20*; *μ=10*^*-6*^.

Finally, we show that the same trends are observed in a spatially explicit agent-based model, although the effect is weaker. A two-dimensional stochastic agent-based model is considered, where reproduction of agents is only possible into the eight nearest neighboring spots (see Materials and Methods for details). As shown in the Supplementary Materials, Section 3, the trends described for the deme-structured model hold true, although fitness is altered to a lesser extent by proportionally scaling birth and death rates.

## Discussion and Conclusion

The life-time reproductive output of individuals in a population is typically considered an indicator for their relative fitness. A larger reproductive output is thought to lead to a selective advantage, while a smaller reproductive output is thought to result in a selective disadvantage. Here we have shown that in structured populations this notion can break down if a mutant with a larger or smaller reproductive output is also characterized by proportionally scaled birth and death rates. Hence, a mutant with a larger reproductive output compared to wild-type can be disadvantageous if it has increased birth and death rates. Similarly, a mutant with a lower reproductive output can become advantageous if it is also characterized by reduced birth and death rates. The notion that a proportionally scaled change in birth and death rates can fundamentally alter fitness properties of a mutant in structured populations revises our understanding of how life-time reproductive success correlates with the Darwinian fitness of a mutant. The observation that this phenomenon occurs in deme-structured population with migration occurring to any deme in the system indicates that the driver of this effect is population fragmentation rather than spatially restricted divisions. We note that in models assuming well-mixed populations, these results are not observed and Darwinian fitness correlates with life-time reproductive success (a deviation from this is observed in small populations and disappears with an increase in *N*, see [17]).

These results not only change our interpretation of selection and evolutionary dynamics, but also increase our understanding of evolution and invasion across many biological and ecological systems. On the biomedical side, tumor progression involves the emergence of cells characterized both by an increased proliferation and death rate [3-9] in spatially structured cell populations. According to our analysis, such cells can in fact be characterized by a selective disadvantage even if their reproductive output is higher than that of the resident cell population, making it more challenging for mutant cells to emerge. A larger increase in reproductive output would be required to overcome the negative consequences of the increased turnover. Even if the cells are selected for, the degree of the selective advantage in the context of higher turnover would be much smaller than anticipated by previous evolutionary theory. Among viruses, faster replication rates can be associated with higher death rates of infected cells or faster decay rates of the virus particles [1, 2, 18], leading to similar evolutionary consequences. Conversely, variants with a fitness cost (such as some drug resistant cell or virus mutants) could have a higher effective fitness than previously thought if the reduction in reproductive output due to the resistance mutation is also accompanied by reduced turnover.

In ecological systems, reproduction and death can also be coupled, such that some species may have proportionally scaled parameters. In many animals, pregnancy can result in reduced mobility and escape velocity during predator attacks, for example in fish [19], dolphins [20], snails [21], snakes [22], birds [23] or lizards [24]. This can lead to an increased death rate in pregnant individuals, which has been demonstrated experimentally for the mosquitofish *Gambusia affinis* [10]. In these cases, our results suggest that the conditions for the selection of new variants / mutants can be more complicated than indicated by existing evolutionary theory, and that relative life-time reproductive success of a new mutant correlates in complex ways with selection. Our results can be further relevant for the dynamics of invasive species, as they spread in new habitats and compete with resident species. For example, for the invasive snail species *Melanoides tuberculata*, it has been found that rapid population growth in the new habitats can be countered by a high mortality rate, due to crayfish-induced predation [21]. Our results help us better understand how this competition plays out, and how this impacts management strategies.

## Materials and Methods

### The deme-structured model

Stochastic Gillespie type simulations were performed to study deme structured dynamics, see also [25]. Consider *D* demes. In each deme, stochastic Gillespie-type simulations are performed, based on the following ordinary differential equations. *dx/dt = r*_*w*_*x[1-(x+y)/K] – d*_*w*_ *x*; *dy/dt = r*_*m*_ *y[1-(x+y)/K] – d*_*m*_*y*. The variables *x* and *y* denote wild-type and mutant populations, respectively. The reproduction and death rates are denoted by *r*_*w*_ and *d*_*w*_ for wild-type individuals and *r*_*m*_ and *d*_*m*_ for mutants, and the carrying capacity is denoted by *K*. We assume *r*_*m*_*=(1+s)* τ *r*_*w*_, *d*_*m*_*=* τ *d*_*w*_, where τ represents the scaling of mutant birth and death rates, and *s* the selection coefficient (*s=0* for quasi-neutral mutants). In addition to the local dynamics, a division event is assumed to lead to offspring placement to a different deme that is randomly chosen from the whole system, with propensities *ε r*_*w*_ *x [1-(x+y)/K]* and *ε r*_*m*_ *y[1-(x+y)/K]*. This migration event is non-spatial and is assumed to be successful only if the population size in the target deme is lower than carrying capacity, *K*. As initial conditions, the wild-type population is set to the equilibrium value in each deme (rounded to the nearest integer). The Gillespie simulation was run for 10,000 time-steps and a single mutant individual was added to a randomly chosen deme if the population size in that deme was below carrying capacity. Otherwise, a new deme was chosen until the mutant was placed successfully. The simulation was run until either the mutant or wild-type population went extinct, and the outcome was recorded.

### Spatial, stochastic agent-based model

We consider a grid that contains *nxn* spots, which may be either empty or contain a wild-type or mutant individual. At each time step, the grid is sampled randomly until *M* spots that contain individuals have been found (where *M* denotes the current number of nonempty spots). If a chosen spot is nonempty, there is a probability for the individual to die (with probabilities *D*_*w*_ and *D*_*m*_*=* τ *D*_*w*_ for wild-types and mutants, respectively), and there is a probability to attempt a reproduction event with probabilities R_w_ and *R*_*m*_*=(1+s)* τ *R*_*w*_, respectively. If an individual is chosen for reproduction, one of the 8 nearest neighboring spots is selected randomly, and if that spot is empty, the offspring individual is placed there (and reproduction does not proceed otherwise). Periodic boundary conditions were assumed. To start the simulation, the average wild-type population size when fluctuating around steady state was determined numerically, and this number of individuals was randomly seeded across the grid. The simulation was run for 500 time-steps to allow the spatial densities to equilibrate, and then a randomly chosen wild-type individual was replaced with a mutant. The simulation was run until either the mutant or wild-type population went extinct, and the outcome was recorded.

## Notes

### Competing Interest Statement

The authors have declared no competing interest.

## References

[1] De Paepe, M. & Taddei, F. 2006 Viruses’ life history: towards a mechanistic basis of a trade-off between survival and reproduction among phages. PLoS biology 4, e193.

[2] García-Villada, L. & Drake, J.W. 2013 Experimental selection reveals a trade-off between fecundity and lifespan in the coliphage Qß. Open biology 3, 130043.

[3] Leoncini, L., Del Vecchio, M.T., Megha, T., Barbini, P., Galieni, P., Pileri, S., Sabani, E., Gherlinzoni, F., Tosi, P. & Kra, R. 1993 Correlations between apoptotic and proliferative indices in malignant non-Hodgkin’s lymphomas. The American journal of pathology 142, 755.

[4] Aihara, M., Truong, L.D., Dunn, J.K., Wheeler, T.M., Scardino, P.T. & Thompson, T.C. 1994 Frequency of apoptotic bodies positively correlates with Gleason grade in prostate cancer. Human pathology 25, 797–801.

[5] Tatebe, S., Ishida, M., Kasagi, N., Tsujitani, S., Kaibara, N. & Ito, H. 1996 Apoptosis occurs more frequently in metastatic foci than in primary lesions of human colorectal carcinomas: analysis by terminal-deoxynucleotidyl-transferase-mediated dUTP-biotin nick end labeling. Interna<onal journal of cancer 65, 173–177.

[6] Vakkala, M., Lähteenmäki, K., Raunio, H., Pääkkö, P. & Soini, Y. 1999 Apoptosis during breast carcinoma progression. Clinical Cancer Research 5, 319–324.

[7] Komaki, R., Fujii, T., Perkins, P., Ro, J.Y., Allen, P.K., Mason, K.A., Mountain, C.F. & Milas, L. 1996 Apoptosis and mitosis as prognostic factors in pathologically staged N1 nonsmall cell lung cancer. Interna<onal Journal of Radia<on Oncology* Biology* Physics 36, 601–605.

[8] O’neill, A., Staunton, M. & Gaffney, E. 1996 Apoptosis occurs independently of bcl-2 and p53 over-expression in non-small cell lung carcinoma. Histopathology 29, 45–50.

[9] Puglisi, F., Minisini, A.M., Aprile, G., Barbone, F., Cataldi, P., Artico, D., Damante, G., Beltrami, C.A. & Di Loreto, C. 2002 Balance between cell division and cell death as predictor of survival in patients with non-small-cell lung cancer. Oncology 63, 76–83.

[10] Laidlaw, C.T., Condon, J.M. & Belk, M.C. 2014 Viability costs of reproduction and behavioral compensation in western mosquitofish (Gambusia affinis). PloS one 9, e110524.

[11] Parsons, T.L., Quince, C. & Plotkin, J.B. 2010 Some consequences of demographic stochasticity in population genetics. Gene<cs 185, 1345–1354.

[12] Parsons, T.L. & Quince, C. 2007 Fixation in haploid populations exhibiting density dependence II: The quasi-neutral case. Theore<cal popula<on biology 72, 468–479.

[13] Bhat, A.S. & Gural, V. 2024 Eco-evolutionary dynamics for finite populations and the noise-induced reversal of selection. bioRxiv, 2024.2002.2019.580940.

[14] Wodarz, D., Goel, A. & Komarova, N.L. 2017 Effect of cell cycle duration on somatic evolutionary dynamics. Evolu<onary applica<ons 10, 1121–1129. (doi:10.1111/eva.12518).

[15] Gillespie, D.T. 1976 General Method for Numerically Simulating Stochastic Time Evolution of Coupled Chemical-Reactions. Journal of Computa<onal Physics 22, 403–434. (doi:Doi 10.1016/0021-9991(76)90041-3).

[16] Wodarz, D. & Komarova, N.L. 2023 Mutant fixation in the presence of a natural enemy. Nature communica<ons 14, 6642.

[17] Kaveh, K., Komarova, N.L. & Kohandel, M. 2015 The duality of spatial death–birth and birth–death processes and limitations of the isothermal theorem. Royal Society open science 2, 140465.

[18] Marianneau, P., Cardona, A., Edelman, L., Deubel, V. & Desprès, P. 1997 Dengue virus replication in human hepatoma cells activates NF-kappaB which in turn induces apoptotic cell death. Journal of virology 71, 3244–3249.

[19] Ghalambor, C.K., Reznick, D.N. & Walker, J.A. 2004 Constraints on adaptive evolution: the functional trade-off between reproduction and fast-start swimming performance in the Trinidadian guppy (Poecilia reticulata). The American naturalist 164, 38–50.

[20] Noren, S.R., Redfern, J.V. & Edwards, E.F. 2011 Pregnancy is a drag: hydrodynamics, kinematics and performance in pre-and post-parturition borlenose dolphins (Tursiops truncatus). Journal of Experimental Biology 214, 4151–4159.

[21] Work, K. & Mills, C. 2013 Rapid population growth countered high mortality in a demographic study of the invasive snail, Melanoides tuberculata (Müller, 1774), in Florida. Aqua<c Invasions 8.

[22] Webb, J.K. 2004 Pregnancy decreases swimming performance of female northern death adders (Acanthophis praelongus). Copeia 2004, 357–363.

[23] Ghalambor, C.K. & Martin, T.E. 2001 Fecundity-survival trade-offs and parental risk-taking in birds. Science 292, 494–497.

[24] Shine, R. 2003 Locomotor speeds of gravid lizards: placing’costs of reproduction’within an ecological context. Func<onal Ecology, 526–533.

[25] Wodarz, D. & Komarova, N.L. 2020 Mutant evolution in spatially structured and fragmented expanding populations. Gene<cs 216, 191–203.

