## Supplementary Materials for "Population structure reverses selection of variants with proportionally scaled birth and death rates"

### Supplemental information

### 1 Probability of mutant fixation in a fragmented population: a coarse-grained approximation

#### 1.1 Set-up and notations

Suppose the dynamics of two species (the wild-type and the mutant) are described by a death and birth process (e.g. a stochastic variant of the Verhulst process [1, 2], such as studied in [3]).

Let us denote by $r_w$ and $d_w$ the w.t. the intrinsic, per capita wild-type division and death rates, and by $r_m$ and $d_m$ the intrinsic, per capita mutant division and death rates. We assume that

$$r_w > d_w, \quad r_m > d_m. \tag{1}$$

Here we will study the probability of mutant fixation under different values of $r_m$ and $d_m$ relative to their wild-type counterparts, but special attention will be paid to the so-called quasi-neutral mutants. We define quasi-neutral mutants as those satisfying

$$r_m = \tau r_w, \quad d_m = \tau d_w. \tag{2}$$

That is, the mutants' rates are proportional to the wild type rates with the factor $\tau$. For $\tau > 1$ the mutants are characterized by a faster turnover, while for $\tau < 1$ they are characterized by a slower turnover.

Here we present some details of calculations focusing on probability of mutant fixation in fragmented and non-fragmented systems, exploring different models of migration and mutant initiation. Tables 1-3 summarize the notations used here.

| Quantity | Notation |
|---|---|
| Division and death rates of wild-type individuals | $r_w, d_w$ |
| Division and death rates of mutant individuals | $r_m, d_m$ |
| Carrying capacity of a Verhulst deme | $K$ |
| Mean population in a deme populated by wild-types and mutants | $N_w$ and $N_m$ |
| Number of demes | $D$ |
| Turnover factor for quasi-neutral mutants | $\tau$ |
| Selection coefficient for mutants | $s$ |

Table 1: System parameters and their notations.

| Quantity | Notation |
|---|---|
| Fitness parameter of a mutant individual (no subscript if constant) | $f_j$ |
| Fitness parameter of a mutant deme (no subscript if constant) | $\mathcal{F}_J$ |

Table 2: Notations for fitness parameters.

| Quantity | Notation |
|---|---|
| Prob. of mut. fix. in a deme, starting with $j$ mut. and $i$ w.t. individuals | $\pi_{ij}$ |
| Prob. of mut. fix. in a single deme, starting with 1 mut. in a population of w.t. individuals, under different models of mutant seeding | $\rho_m$ |
| Prob. of w.t. fix. in a single deme, starting with 1 w.t. in a population of mutants, under different models of wild-type seeding | $\rho_w$ |
| Prob. of a mutant deme fixation starting from a single mutant deme (in the coarse-grained model) | $P_{deme}$ |
| Prob. of mutant fixation in a fragmented multi-deme system, starting with 1 mutant individual | $\Pi_1^{frag}$ |
| Prob. of mutant fixation in an equivalent non-fragmented system, starting with 1 mutant individual | $\Pi_1^{non}$ |

Table 3: Notations for fixation probabilities.

#### 1.2 Probability of mutant fixation in a single deme

##### 1.2.1 Model formulation and the quasi-stationary probability distribution

Consider a death and birth process where the deaths and the births are decoupled, resulting in demographic fluctuations in the population size. Suppose that parameter $K$ (the carrying capacity) is fixed. Denote the states of the system by a pair of integers $(i, j)$ where $i$ is the number of wild-type individuals and $j$ is the number of mutants. In a version of the Verhulst process, we assume that the following processes can occur during an infinitesimal time-interval, $\Delta t$:

$$P_{(i,j)\to(i+1,j)} = ir_w \left[1 - \frac{i+j}{K}\right]_+ \Delta t, \tag{3}$$

$$P_{(i,j)\to(i,j+1)} = jr_m \left[1 - \frac{i+j}{K}\right]_+ \Delta t, \tag{4}$$

$$P_{(i,j)\to(i-1,j)} = id_w \Delta t, \tag{5}$$

$$P_{(i,j)\to(i,j-1)} = jd_m \Delta t, \tag{6}$$

$$P_{(i,j)\to(i,j)} = 1 - \left((ir_w + jr_m)\left[1 - \frac{i+j}{K}\right]_+ (id_w + jd_m)\right)\Delta t, \tag{7}$$

with all the other transition probabilities assumed to be zero. Here $[z]_+$ denotes the positive part,

$$[z]_+ = \begin{cases} z, & z \geq 0, \\ 0, & z < 0. \end{cases}$$

This process has only one absorbing state, $(i, j) = (0, 0)$, which is the total population extinction. For our purposes, however, we will assume that extinction takes a very long time. Intuitively, provided that the death rates are not too close to the division rates, $K$ is sufficiently large, and the initial condition is around $N = K(1 - d_w/r_w)$, we can ignore the possibility of population extinction on the time-scales of interest (see [3, 4] for the estimate of the extinction time given the kinetic parameters).

**Quasi-stationary probability distribution.** To study the system, it is useful to calculate the quasi-stationary probability distribution of the deme size (see also [3]). In a deme with wild-type individuals only, denote the probability of finding the system in state $i$ as $P_i$. We have

$$d_w P_1 = 0, \tag{8}$$

$$d_w P_2 2 - P_1 \left( r_w \left( 1 - \frac{1}{K} \right) + d_w \right) = 0, \tag{9}$$

$$r_w P_{i-1}(i-1)\left(1 - \frac{i-1}{K}\right) + d_w P_{i+1}(i+1) - P_i i \left( r_w \left(1 - \frac{i}{K}\right) + d_w \right) = 0, \quad 1 < i < K, \tag{10}$$

$$r_w P_{K-1}(K-1)\left(1 - \frac{K-1}{K}\right) - P_K K d_w = 0. \tag{11}$$

It follows that all the probabilities except $P_0$ are zero. Therefore, in order to obtain the long-term distribution in the absence of extinction, we remove the transition from state 1 to state 0, such that equation (9) for $i = 1$ becomes

$$d_w P_2 2 - P_1 r_w \left(1 - \frac{1}{K}\right) = 0, \tag{12}$$

and equation (8) becomes an identity. System (12,10,11) can be solved together with the normalization $\sum_{i=1}^{K} P_i = 1$ and results in a stationary probability distribution that correctly describes results of numerical simulations, see figure S1. Note that the operation of cutting the transition to zero is equivalent to the operation of adding a flux from state 0 to state 1 in this context and leads to the same solution for $P_1, \ldots, P_K$.

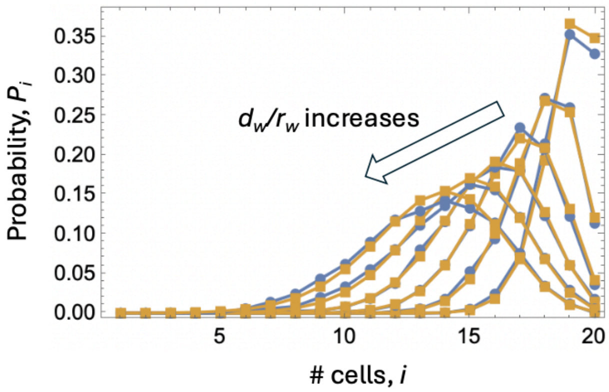

Figure S1: Stationary probability distribution calculated by solving system (10,11,9) (yellow) and obtained from running stochastic simulations (blue). Results for $d_w/d_r = 0.1, 0.2, \ldots, 0.6$ are shown; $K = 20$.

**Probability for a deme to be non-full.** A useful quantity related to this calculation is the probability of a deme to be non-full (we will use this quantity when studying migration of individuals from deme to deme). Denote by $P_w^{non-full}$ the probability that the population of a deme at a quasi-equilibrium state is less than $K$. We have

$$P_w^{non-full} = 1 - P_K, \tag{13}$$

where $P_K$ is a component of the normalized solution of system (12,10,11). The probability for a deme to be non-full increases with the death-to-division rate and the carrying capacity, and can be approximated by

$$P_w^{non-full} \approx 1 - \exp\left(-\frac{K^2}{K-1}\frac{d_w}{r_w}\right), \tag{14}$$

as shown in figure S2. In particular, two limits are of interest:

- *High-density limit.* If $d_w/r_w \ll 1/K$ (we neglected 1 compared to $K$ in this condition), we have
$$P_w^{non-full} \approx K\frac{d_w}{r_w}. \tag{15}$$
- *Low-density limit.* If $d_w/r_w \gg 1/K$, we have
$$P_w^{non-full} \approx 1. \tag{16}$$

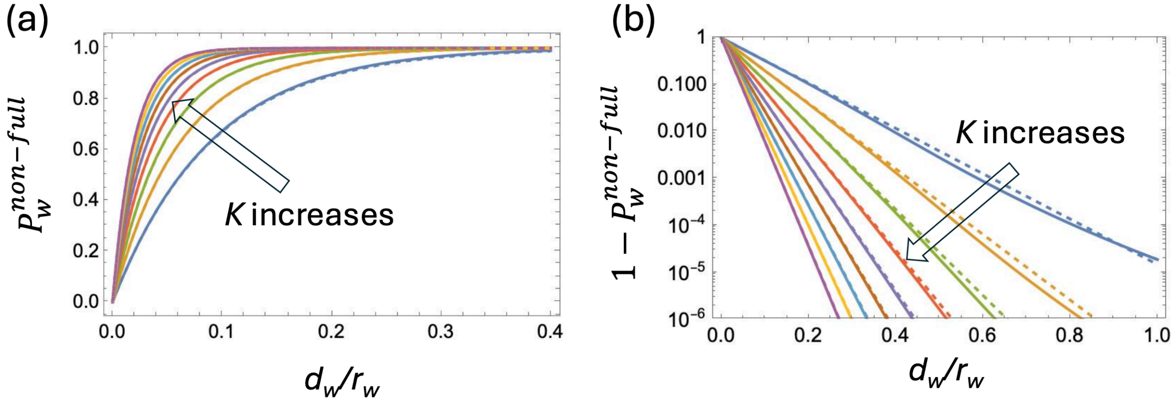

Figure S2: The probability for a deme to be non-full. (a) $P_w^{non-full}$ plotted as a function of the death-to-division ratio, $d_w/r_w$. (b) $1 - P_w^{non-full}$ vs $d_w/r_w$, plotted on a logarithmic scale. The different colors correspond to different values of the carrying capacity parameter, which was taken to be $10, 15, 20, \ldots, 50$. Solid lines represent formula (13), obtained by explicitly solving system (12,10,11). Dashed lines are the approximation (14).

##### 1.2.2 Calculating mutant fixation probability in a deme

To study the probability of mutant fixation in a deme, let us artificially stop the process once one of the boundary states is reached, that is, we will make all the states $(i, 0)$ with $i \geq 0$ and all the states $(0, j)$ with $j > 0$ absorbing. Then, we will say that "mutants reached fixation" if one of the states $(0, j)$ with $j > 0$ is reached. In the context of the original process, this simply means that we are interested in the probability of the wild type individuals going extinct first (with eventually the whole population going extinct).

Denote by $\pi_{ij}$ the probability of mutant fixation (with the above definition) starting from $i$ wild type and $j$ mutant individuals. This quantity satisfies the following equations:

$$ir_w\left(1-\frac{i+j}{K}\right)\pi_{i+1,j}+jr_m\left(1-\frac{i+j}{K}\right)\pi_{i,j+1}+id_w\pi_{i-1,j}+jd_m\pi_{i,j-1}$$
$$-\left((ir_w+jr_m)\left(1-\frac{i+j}{K}\right)+(id_w+jd_m)\right)\pi_{ij}=0,\quad i>0,j>0,i+j<K, \tag{17}$$
$$id_w\pi_{i-1,j}+jd_m\pi_{i,j-1}-(id_w+jd_m)\pi_{ij}=0,\quad i+j\geq K \tag{18}$$

with the boundary conditions given by

$$\pi_{k,0}=0,\quad \pi_{0,k}=1,\quad k>0. \tag{19}$$

System (17-19) can be solved recursively up to an arbitrary value of $i+j=K_0\geq K$. In particular, of interest is the probability of mutant fixation starting from its quasi-stationary state, where the population is approximately at its deterministic equilibrium,

$$N_w=K(1-d_w/r_w). \tag{20}$$

Denote by $\rho_m$ the probability of mutant fixation starting from a single mutant in a population of wild-type individuals, under the process with demographic fluctuations, equations (3-7). We have by solving system (17-19),

$$\rho_m=\pi_{N_w-1,1}. \tag{21}$$

In this study we will also need to calculate the probability of a wild-type fixation in a population of mutants, which we denote $\rho_w$. The quasi-stationary state in a mutant deme is given by

$$N_m=K(1-d_m/r_m), \tag{22}$$

and we have

$$\rho_w=\pi_{N_m-1,1}, \tag{23}$$

where we use the solution of system (17-19), where the wild-type coefficients were replaced by the mutant coefficients, $r_w\leftrightarrow r_m$ and $d_w\leftrightarrow d_m$.

When discussing the probability of mutant fixation in a small deme with a fluctuating size, it is important to correctly capture the process of mutant initiation. Suppose that a deme population has reached a quasi-equilibrium state, where it oscillates around its equilibrium value. We further suppose that a mutant migrates from the outside to the deme. Then the probability of this mutant to fixate in the population is given, instead of equation (21), by

$$\rho_m=\sum_{i=1}^{K}P_i\pi_{i,1}, \tag{24}$$

where the values $\pi_{i,1}$ are obtained by solving system (17-19). The expression for $\rho_w$ is also given by the right hand side of equation (24), except now the values $\pi_{i,1}$ are solutions of system (17-19), where we swap the mutant and wild-type kinetic rates: $r_w\leftrightarrow r_m$ and $d_w\leftrightarrow d_m$.

##### 1.2.3 Fixation probability of quasi-neutral mutants: comparison with the diffusion approximation

Let us focus on quasi-neutral mutants, equation (2). Note that for such mutants, $N_w = N_m = N$ (equations (22,20)).

Approximate methods can be applied to obtain the probability of mutant fixation, if the population size is large. In [5, 6], the diffusion approximation for quasi-neutral mutants was used to solve a similar system, with the difference that instead of density-controlled divisions, density-controlled death was implemented. The approximate solution for the probability of mutant fixation obtained for this system is given by $\pi_{N-1,1} \approx \rho_m^{diff}$, where

$$\rho_m^{diff} = \frac{1}{N}\frac{2}{\tau+1} \tag{25}$$

Numerical simulations confirm that this formula works well for both the process with density controlled death and the process with density controlled divisions, equations (3-7), as long as the population size is large, see S3(a) and the text below. Remarkably, this is the same expression as was obtained for the probability of quasi-neutral mutant fixation in the (constant-population) death-birth Moran process [7].

Here we are interested in the deviation of the probability of mutant and wild-type fixation (equations (21,23)) from the diffusion approximation. The latter, for the probability of mutant fixation, is given by equation (25), and the corresponding quantity for wild-type fixation under the diffusion approximation is given by

$$\rho_w^{diff} = \frac{1}{N}\frac{2}{1/\tau+1}.$$

As mentioned above, for large populations when the diffusion approximation holds, we have $\rho_m \to \rho_m^{diff}$ and $\rho_w \to \rho_w^{diff}$. For smaller populations, however, diffusion approximation breaks down.

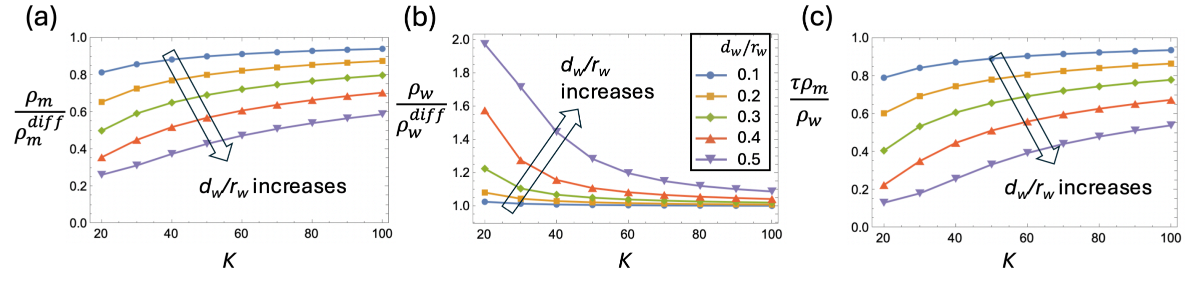

Figure S3: Probabilities of fixation in a population with demographic fluctuations (system (17-19)), compared to their values obtained by diffusion approximation, in a system with quasi-neutral mutants. (a) The ratio $\rho_m/\rho_m^{diff}$, as a function of the carrying capacity, $K$, plotted for several values of the $d_w/r_w$ ratio. For each point, the population size was taken $N = K(1 - d_w/r_w)$, which is an integer number for all the cases. (b) The same for the ratio $\rho_w/\rho_w^{diff}$. (c) The same for the quantity $\tau\rho_m/\rho_w$. The ratios $d_w/r_w$ are shown in the inset in panel (b). We used $\tau = 10$ in all panels.

Figure S3(a) shows the ratio of the mutant fixation probability in a population with demographic fluctuations (system (17-19)) and the corresponding value obtained by diffusion approximation, $\rho_m/\rho_m^{diff}$. We can see that for $\tau = 10 > 1$, which was used in this figure, this ratio is smaller than 1, and tends to one when the carrying capacity ($K$) increases and the death-to-division ratio decreases. Figure S3(b) shows a similar plot for the probability of wild-type fixation. The ratio is generally greater than one ($1/\tau = 1/10 < 1$), and again it approaches when the carrying capacity increases and the death-to-division ratio decreases. Panel (c) shows quantity

$$\mathcal{F} = \tau\rho_m/\rho_w,$$

which should be compared with the corresponding quantity under diffusion approximation,

$$\tau\rho_m^{diff}/\rho_w^{diff} = 1.$$

We can see that, for fast mutants ($\tau > 1$) in fluctuating populations, the probability of mutant fixation is smaller than that calculated under diffusion approximation, and the probability of mild-type fixation is larger than that calculated under diffusion approximation. As a result, the ratio $\rho_m/\rho_w$ is smaller than $1/\tau$.

#### 1.3 Probability of mutant fixation in a fragmented system

##### 1.3.1 The coarse-grained approximation

Next, we consider a system of $D$ demes where wild-type and mutant individuals can migrate between demes. We assume that a migrating individual is equally likely to land on any deme, in other words, there is no spatial structure in this model. The key assumption is that the migration rate is low compared to the processes within demes, such that typically, after a migration event, the deme populations will become homogeneous (through fixation or extinction of the immigrant type), before the next migration event happens. We will use the coarse grained approximation (see [8]) to calculate the probability of mutant fixation. The key idea is to approximate the probability of mutant fixation, $\Pi_1^{frag}$, as

$$\Pi_1^{frag} = \rho_m P_{deme}, \tag{26}$$

where the probability of a mutant deme fixation (starting from a single mutant deme) is denoted by $P_{deme}$, and the probability of mutant fixation in a single deme is $\rho_m$, see table 3.

To calculate the probability of deme fixation, $P_{deme}$, consider the population of $D$ demes (populated entirely either by mutant or by wild-type individuals). We denote by $J$ the number of mutant demes; the number of wild-type demes in $D - J$. Let us denote the rate, at which the number of mutant demes increases by one, by $W_J^+$, and the rate, at which the number of mutant demes decreases by one ($W_J^-$). Denote by $h_J$ the probability of mutant deme fixation starting from $J$ mutant demes. We have, by the standard one-step analysis,

$$0 = W_J^+ h_{J+1} + W_J^- h_{J-1} - h_J(W_J^+ + W_J^-), \quad h_0 = 0, \quad h_D = 1. \tag{27}$$

Denoting

$$\mathcal{F}_J = \frac{W_J^+}{W_J^-},$$

equation (27) can be rewritten as

$$0 = \mathcal{F}_J h_{J+1} + h_{J-1} - h_J(\mathcal{F}_J + 1), \quad h_0 = 0, \quad h_D = 1. \tag{28}$$

The next steps are to construct the expressions for $W_J^+$, $W_J^-$, and $\mathcal{F}_J$, solve equation (28), and determine

$$P_{deme} = h_1. \tag{29}$$

##### 1.3.2 Deme fitness for different migration models

For migrations, a number of modeling assumptions are possible. The first choice to make is the mechanism by which migration is carried out. Here we distinguish two major types of migration models:

(1) **Reproduction-independent migration.** We can assume that migration happens as a separate event (distinct from birth and death events), and the migration rate is either the same for both types, or that it is proportional to division rate of individuals.

(2) **Migration by division.** Alternatively, we could assume that migration is incorporated in divisions, when with a given small probability ($\epsilon$) the offspring of a divided individual is transported to a different deme.

As before, we assume that the migration rate is low, such that the demes are homogeneous most of the time.

The second modeling choice concerns the migration event destination. Two possibilities are considered here:

(a) **Crowd-controlled migration:** Migration can only occur to a deme whose population size is smaller than $K$. If the target deme is full (size $K$) then the migration event is aborted.

(b) **Crowd-independent migration:** Migration happens regardless of whether or not a deme is full.

Below we calculate the rate, at which the number of mutant demes increases by one ($W_J^+$) and decreases by one ($W_J^-$), and the resulting expression for the mutant deme fitness. Despite this variety of modeling choices, as we show, the resulting expression for the deme fitness is the same or similar in many cases.

We will begin by assuming that the migration process is **reproduction-independent** (choice (1) above) and **crowd-controlled** (choice (a) above). Consider the population of $J$ mutant demes and $D_J$ wild-type demes. The populations of the demes fluctuate around their equilibrium values. Wild type and mutant individuals can migrate between demes. A change in the number of mutant and wild-type demes happens through the process of deme conversion. The rate of conversion is given by the rate at which an individual of a given type lands on a deme of a different type, times the probability of fixation of the migrant. As before, denote by $N_w$ and $N_m$ the equilibrium value of the wild-type and mutant demes, see equations (20,22). We have, for the rates of increase and decrease in the number of mutant demes,

$$W_J^+ \propto JN_m\mu_m \times \frac{D-J}{D} \times P_w^{non-full} \times \rho_m, \tag{30}$$

$$W_J^- \propto (D-J)N_w\mu_w \times \frac{J}{D} \times P_m^{non-full} \times \rho_w, \tag{31}$$

where the right hand side of the expression for $W_J^+$ is proportional to the propensity of mutants to migrate (which is the total number of mutants, $JN_m$, times the migration rate, $\mu_m$), land on a wild-type deme ($\frac{D-J}{D}$), find that the chosen wild-type deme is non-full ($P_w^{non-full}$), and get fixated there ($\rho_m$). The expression for $W_J^-$ is constructed similarly. The probability that a wild-type deme is non-full, $P_w^{non-full}$, is given by equation (13). The probability that a mutant deme is non-full, $P_m^{non-full}$, requires switching the parameters $r_w, d_w$ in system (12,10,11) to parameters $r_m, d_m$.

From these equations, the fitness parameter for mutant demes is given by

$$\mathcal{F}_J = \frac{W_J^+}{W_J^-} = \frac{N_m P_w^{non-full}}{N_w P_m^{non-full}} \frac{\mu_m \rho_m}{\mu_w \rho_w} = \mathcal{F}, \tag{32}$$

which is independent of the number of mutant demes, $J$.

Different assumptions on the migration process will modify the expressions in the following way. For **migration by division** (choice (2) above), the rate of migration is $\epsilon d_m$ for mutants and $\epsilon d_w$ for wild-type individuals (because at the equilibrium, the rate of successful divisions is equal to the rate of death). These expressions will replace the rates $\mu_m$ and $\mu_w$, respectively, in formulas (30) and (31). For **crowd-independent migration** (choice (b) above), the rate of migration does not depend on whether or not the target deme is full. Therefore, expressions $P_m^{non-full}$ and $P_w^{non-full}$ should be replaced by 1 in formulas (30) and (31).

| | (1) Reproduction-independent migration | (2) Migration by division |
|---|---|---|
| (a) Crowd-controlled migration | $\frac{N_m P_w^{non-full}}{N_w P_m^{non-full}} \frac{\mu_m \rho_m}{\mu_w \rho_w}$ | $\frac{N_m P_w^{non-full}}{N_w P_m^{non-full}} \frac{d_m \rho_m}{d_w \rho_w}$ |
| (b) Crowd-independent migration | $\frac{N_m}{N_w} \frac{\mu_m \rho_m}{\mu_w \rho_w}$ | $\frac{N_m}{N_w} \frac{d_m \rho_m}{d_w \rho_w}$ |

Table 4: Expressions for the mutant deme fitness, $\mathcal{F}$, for different assumptions on the migration process.

Table 4 lists all the combinations of the assumptions. Note that the expressions presented in the table are quite general, in the sense that the microscopic assumptions on the within-deme dynamics can vary. In the model implemented here (that is, a Verhulst process with density-dependent divisions), the expected population sizes of demes are given by equations (20,22), the probability to be non-full is equation (13), obtained by solving system (12,10,11), and the probability of fixation is given by equation (24). Similar expressions can be derived for a more general birth-death process in the demes, where per-individual division and death rates are some functions of the numbers of wild-type and mutant individuals. The present theory requires that the demes' populations fluctuate around a finite quasi-equilibrium size, $N_w$ and $N_m$, and (for the case of migration by division) that the two species have an intrinsic division rate (what we called $r_w, d_w$).

##### 1.3.3 Special cases: high-density mode, low-density mode, and quasi-neutral mutants

**High-density mode.** As follows from section 1.2.1, we can use simple approximations for the quantities $P_w^{non-full}$ and $P_m^{non-full}$. In particular, if the death-to-division ratios satisfy $\frac{d_w}{r_w} \ll \frac{1}{K}$, $\frac{d_m}{r_m} \ll \frac{1}{K}$, the demes are characterized by high density and are at capacity most of the time. Therefore, the equilibrium deme sizes satisfy $N_m \approx N_w \approx K$. Further, using approximation (15) and a similar approximation for $P_m^{non-full}$ in the expressions of deme fitness, we have $P_w^{non-full} \approx K\frac{d_w}{r_w}$, $P_m^{non-full} \approx K\frac{d_m}{r_m}$, and instead of table 4, we have simplified expressions for mutant deme fitness, that are summarized in table 5.

| | (1) Reproduction-independent migration | (2) Migration by division |
|---|---|---|
| (a) Crowd-controlled migration | $\frac{d_w r_m}{r_w d_m} \frac{\mu_m \rho_m}{\mu_w \rho_w}$ | $\frac{d_w r_m}{r_w d_m} \frac{d_m \rho_m}{d_w \rho_w}$ |
| (b) Crowd-independent migration | $\frac{\mu_m \rho_m}{\mu_w \rho_w}$ | $\frac{d_m \rho_m}{d_w \rho_w}$ |

Table 5: High-density mode, where $\frac{d_w}{r_w} \ll \frac{1}{K}$, $\frac{d_m}{r_m} \ll \frac{1}{K}$. Expressions for the mutant deme fitness, $\mathcal{F}$, are given for different assumptions on the migration process.

**Low-density mode.** In the opposite limit, where the death-to-division ratios satisfy $\frac{d_w}{r_w} \gg \frac{1}{K}$, $\frac{d_m}{r_m} \gg \frac{1}{K}$, the demes are characterized by low density and rarely reach the carrying capacity. As a consequence, using approximation (16) and a similar approximation for $P_m^{non-full}$ in the expressions of deme fitness, we obtain:

$$\text{(low-density:)} \quad \mathcal{F} = \begin{cases} \frac{N_m}{N_w}\frac{\mu_m \rho_m}{\mu_w \rho_w}, & \text{reproduction-independent migration,} \\ \frac{N_m}{N_w}\frac{d_m \rho_m}{d_w \rho_w}, & \text{migration by division.} \end{cases}$$

In other words, since the demes are almost never full, there is no difference between crowd-controlled and crowd-independent migration models in this case.

**Mutant deme fitness for quasi-neutral mutants.** The expressions for mutant deme fitness (regardless of the population density) simplify significantly in the case of quasi-neutral mutants. Since the equilibrium deme sizes (equations (20,22)) and the solution of system (12,10,11) only depend on the ratio, $d_w/r_w$, for quasi-neutral mutants we have

$$\text{(quasi-neutral:)} \quad N_m = N_w, \quad P_m^{non-full} = P_w^{non-full}.$$

Therefore, for the deme fitness, we obtain,

$$\text{(quasi-neutral:)} \quad \mathcal{F} = \begin{cases} \frac{\mu_m \rho_m}{\mu_w \rho_w}, & \text{reproduction-independent migration,} \\ \tau \frac{\rho_m}{\rho_w}, & \text{migration by division.} \end{cases}$$

If the migration rate in the reproduction-independent migration is taken to be proportional to the division rate, $\mu_m/\mu_w = r_m/r_w$, then the two cases coincide, yielding $\mathcal{F} = \tau\frac{\rho_m}{\rho_w}$. This quantity was studied in Section 1.2.3, see e.g. figure S3(c).

##### 1.3.4 Probability of deme fixation and different initiation models

To calculate the probability of mutant fixation in a fragmented population by using coarse-grain approximation, equation (26), we need to calculate the probability of mutant deme fixation, $P_{deme}$. Since the fitness of a deme, $\mathcal{F}$, is a quantity independent of $J$, solution of equation (28) is given by $h_J = \frac{1-1/\mathcal{F}^J}{1-1/\mathcal{F}^D}$, and therefore from (29) we have

$$P_{deme} = \frac{1 - 1/\mathcal{F}}{1 - 1/\mathcal{F}^D}. \tag{33}$$

Apart from using different migration models, there is also a choice of ways in which the system can be initialized. A more general calculation of the probability of quasi-neutral mutant fixation in a fragmented population with demographic fluctuations will depend on the exact assumptions on both the migration process and the initial mutant placement. We consider several cases here, all of which are described by a slight modification of equation (26), where we used equation (33):

$$\Pi_1^{frag} = \rho_m^{init} \frac{1 - 1/\mathcal{F}}{1 - 1/\mathcal{F}^D}.$$

Here $\rho_m^{init}$ stands for the probability of mutant fixation in the deme where it was initially placed, and the mutant deme fitness is calculated in sections 1.3.2 and 1.3.3. The deme fitness calculations involve quantities $\rho_m$ and $\rho_w$, which are probabilities of mutant and wild-type fixation upon migration, with $\rho_m$ that may or may not be different from $\rho_m^{init}$.

We start by describing two cases for $\rho_m$ and $\rho_w$, which depend on migration assumptions:

- Crowd independent migration. In this case, as derived previously, $\rho_m$ is given by formula (24), where the values $\pi_{i,1}$ are obtained by solving system (17-19). To calculate $\rho_w$, we use the same formula, but when solving system (17-19), we exchange $r_w \leftrightarrow r_m$ and $d_w \leftrightarrow d_m$.

- Crowd controlled migration. Since the migration events that target a deme with a population at or above carrying capacity are canceled, we have

$$\rho_m = \frac{\sum_{i=1}^{K-1} P_i \pi_{i,1}}{\sum_{i=1}^{K-1} P_i},$$

  where again $\pi_{i,1}$ are obtained from system (17-19), and the calculation adjusted for $\rho_w$ in the same way as before.

The initial probability of fixation, $\rho_m^{init}$, depends on the way the first mutant is placed. In the literature, several models are used, see e.g. [9]:

(i) Uniform (by deme) mutant initialization, where one individual in a uniformly randomly chosen wild-type deme is turned into a mutant. In this case, we have

$$\rho_m^{init} = \sum_{i=1}^{K} P_i \pi_{i-1,1}.$$

(ii) Uniform (by individual) mutant initialization, where a uniformly randomly chosen wild-type individual is turned into a mutant. In this case, we have

$$\rho_m^{init} = \frac{\sum_{i=1}^{K} P_i i \pi_{i-1,1}}{\sum_{i=1}^{K} P_i i}.$$

(iii) Temperature mutant initialization, where the initial mutant is assumed to be produced by a dividing individual. We have

$$\rho_m^{init} = \frac{\sum_{i=1}^{K-1} P_i i(1-i/K)\pi_{i,1}}{\sum_{i=1}^{K-1} P_i i(1-i/K)}.$$

(iv) A mutant is introduced by an external migration event. In this case, we simply set $\rho_m^{init} = \rho_m$, where the latter quantity is calculated as described above, in accordance with the migration model used.

###### 1.3.5 Probability of quasi-neutral mutant fixation in fragmented populations and their effective fitness

In Fig. 1 of the main paper, as well as in the examples below, we assume that migration by division is crowd-controlled (model (2a) of section 1.3.2), and that the initialization happens according to scenario (iv) of section 1.3.4.

Figure S4 is similar to Fig. 1(A-C) of the main paper, as it shows the probability of quasi-neutral mutant fixation in a fragmented system as a function of (a) the total population size, by increasing the number of demes, (b) the turnover factor, $\tau$, and (c) the death-to-division ratio, $d_w/dr_w$. The difference compared to the main text figure is that here we explore mutants that experience a slowing in their turnover, that is, $\tau < 1$. The values $\Pi_1^{frag}$ are calculated as explained above (see the solid lines). They are compared with the corresponding functions for the non-fragmented populations, $\Pi_1^{non} = \frac{1}{ND}\frac{1}{\tau+1}$ (the dashed lines). We can see that the fixation probability in a fragmented system is always larger than that that for the non-fragmented population.

We also observe a qualitative difference between the behavior of $\Pi_1^{frag}$ and $\Pi_1^{non}$, as we increase the number of demes, $D$ (and thus the total population size, $N_{tot} = DN$. The probability of quasi-neutral mutant in a non-fragmented population, $\Pi_1^{non}$, is always inversely proportional to the total population size. On the other hand, $\Pi_1^{frag}$ decreases

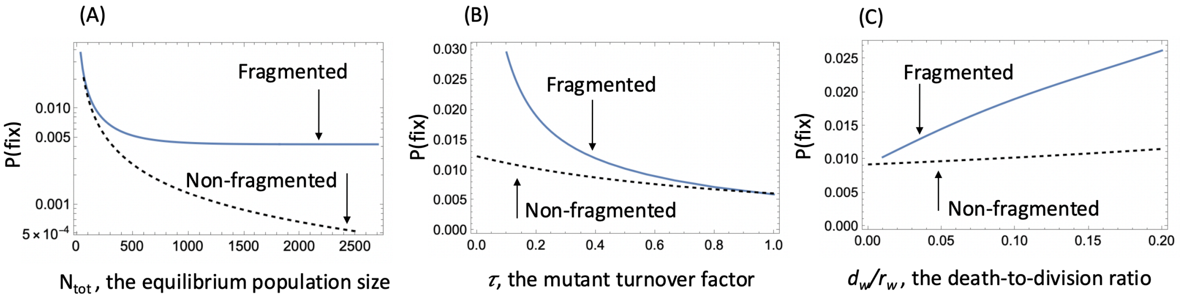

Figure S4: Probability of quasi-neutral mutant fixation probability in the deme-structured model, obtained by the coarse-grained theory, with $\tau < 1$. The solid line is the quantity $\Pi_1^{frag}$, and the dashed line is $\Pi_1^{non}$. (A) The mutant fixation probability increases and reaches a saturation as a function of the total quasi-equilibrium population size, $N_{tot} = ND$, a characteristic of an advantageous mutant. (B) The mutant fixation probability declines with the amount by which the birth and death rates are proportionally increased, $\tau$. (C) The increase of the mutant fixation probability in the deme-structured compared to the non-fragmented population becomes more substantial with higher death to birth ratios. Base parameters were chosen as follows: For (A) $\tau = 0.5$; $D$ varies from 2 to 150. For (B) $D = 9$. For (C) $\tau = 0.2, D = 9$. The rest of the parameters are $d_w/r_w = 0.1; \epsilon = 10^{-4}; K = 20; s = 0$.

exponentially with $N_{tot}$ if $\tau > 1$ (see Fig. 1(A) of the main paper), and it increases and reaches a constant value for $\tau < 1$ (see figure S4(a)). This behavior of the quantity $\Pi_1^{frag}$ suggests that the quasi-neutral mutant in a fragmented populations acquires features of a truly disadvantageous mutant with $\tau > 1$ and a truly advantageous mutant with $\tau < 1$. This can be shown by rewriting the expression for $\Pi_1^{frag}$ as

$$\Pi_1^{frag} = \rho_m^{init} \frac{1 - 1/\mathcal{F}}{1 - 1/\mathcal{F}^D} = \alpha \frac{1 - 1/f_e}{1 - 1/f_e^{N_{tot}}}, \tag{34}$$

where $N_{tot} = ND$, the "effective" fitness parameters of quasi-neutral individuals, $f_e$, is given by

$$f_e = \mathcal{F}^{1/N}, \tag{35}$$

and the proportionality constant, $\alpha$, is independent on the number of demes and is defined as

$$\alpha = \rho_m^{init} \left( \frac{1 - 1/f_e}{1 - 1/f_e^N} \right)^{-1}.$$

The latter expression is the ratio between the probability of the initial mutant to fixate in a single deme, and the probability of a mutant with fitness $f_e$ to fixate in a deme of the same size, if it is subject to the usual Moran dynamics.

Let us consider quasi-neutral mutants in relatively small demes, where deme fitness is not equal to one. If $\tau > 1$, then both $\mathcal{F} < 1$ and $f_e < 1$, then we can see from equation (34) that the fixation probability decays exponentially with the population size, $N_{tot}$. If on the other hand, $\tau < 1$, then $\mathcal{F} > 1$ and $f_e > 1$, then from equation (34), the fixation probability increases and saturates at the level $\alpha \frac{f_e - 1}{f_e}$ as $N_{tot}$ increases. Finally, if $\mathcal{F} = 1$ (which happens if the deme size is large), we have $f_e = 1$, $\rho_m^{init} = \frac{1}{N} \frac{2}{\tau+1}$, and $\alpha = \frac{2}{\tau+1}$, and $\Pi_1^{frag} = \frac{2}{\tau+1} \frac{1}{N_{tot}}$.

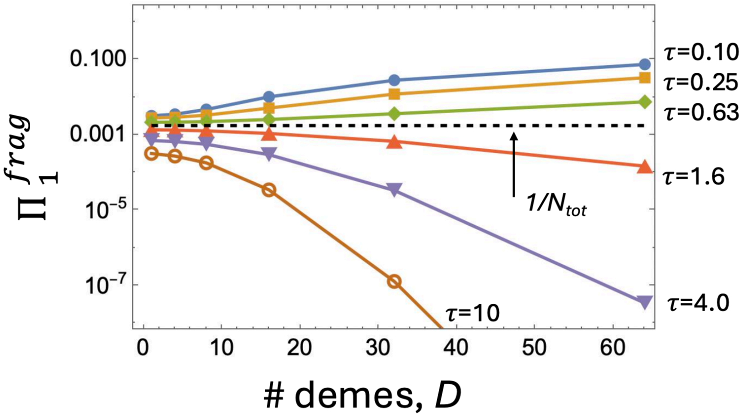

Figure S5: The role of fragmentation in quasi-neutral mutant fixation probability. The quantity $\Pi_1^{frag}$ is plotted as a function of the number of demes, $D$, for a fixed total carrying capacity, $K_{tot} = 640$. Each curve corresponds to a different turnover parameter $\tau dr$ that ranges from $\tau = 10$ to $R = 0.1$, as marked. The probability of neutral mutant fixation in the whole population is shown by the horizontal line. The death-to-division ratio is $d_w/r_w = 0.1$.

To summarize, the behavior of quasi-neutral mutants in fragmented populations with small demes of a fixed size, resembles that of non-neutral mutants with effective fitness (35) and probability of fixation that depends on the total population size as in equation (34), with a constant factor $\alpha$ independent on the deme number.

##### 1.3.6 The role of fragmentation in quasi-neutral and non-neutral mutant fixation in fragmented populations

Next, we examine the influence of fragmentation on the probability of quasi-neutral mutant fixation, under a constant total populatuon size. In Figure S5 we by split the total carrying capacity, $K_{tot}$, into 4,8,16,32, and 64 parts. Six different values of the turnover parameter, $\tau$ were used. Three of them are greater than one; for those parameters, fragmentation results in a decrease of fixation probability. For $\tau = 10$, when the total population is split into 64 demes, the reduction in fixation probability is enormous (about $10^{-10}$). But even for modest values of $\tau$ such as $\tau = 1.6$, the reduction in fixation probability reaches about 10-fold for $D = 64$.

Figure S6 explores the role of fragmentation in fixation dynamics of non-neutral mutants. Two scenarios are presented. In panel (a), the mutants have a division-to-death ratio greater than that of the wild-type (with the selection coefficient $s = 0.01$, which makes them advantageous in a non-fragmented population), but their turnover rate is faster than that for the wild-type. As before, the total carrying capacity is kept constant, and the population is subdivided into 4,8,16,32, and 64 parts. As the degree of fragmentation increases, the probability of mutant fixation decreases and becomes less than that for neutral mutants. In other words, in a subdivided population, a faster turnover rate may turn an advantageous mutant into a disadvantageous mutant.

In panel (b) we consider the opposite scenario, where the mutants have a division-to-death ratio smaller than that of the wild-type (with the selection coefficient $s = -0.01$, which makes them disadvantageous in a non-fragmented population), but their turnover rate is slower than that for the wild-type. As the degree of fragmentation increases, the probability of mutant fixation increases and becomes larger than that for neutral mutants. Therefore, in a subdivided population, a slower turnover rate may turn a disadvantageous mutant into an advantageous mutant.

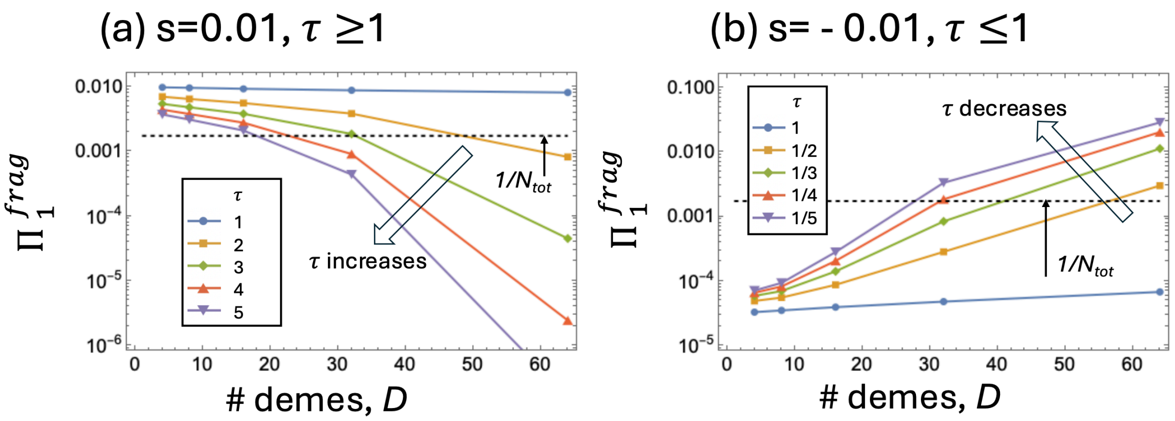

Figure S6: The role of fragmentation in non-neutral mutant fixation probability. (a) Advantageous fast mutants: $s > 0, \tau > 1$. (b) Disadvantageous slow mutants: $s < 0, \tau < 1$. The quantity $\Pi_1^{frag}$ is plotted as a function of the number of demes, $D$, for a fixed total carrying capacity, $K_{tot} = 640$. The probability of neutral mutant fixation in the whole population is shown by the horizontal line. Different curves correspond to different turnover parameters, as indicated in the legend. The rest of the parameters are $r_w = 0.5, r_m = (1+s)\tau r_w, d_w = 0.05, d_m = \tau d_w$.

### 2 Dynamics in the presence of *de novo* mutations

#### 2.1 Selection-mutation balance in fragmented populations

Here we adapt the coarse-grained approach to study the process of *de-novo* mutations in the system. Again we will assume that migration by division is crowd-controlled (model (2a) of section 1.3.2). The calculations can be easily modified t incorporate the other assumptions discussed in the previous section.

As before, denote the number of mutant demes by $J$. In the presence of *de-novo* mutations, the probabilities to increase and decrease the number of mutant demes are given by

$$W_J^+ = JN_m\epsilon d_m \frac{D-J}{D} P_w^{non-full}\rho_m + (D-J)N_w d_w u \rho_m, \quad (36)$$

$$W_J^- = (D-J)N_w\epsilon d_w \frac{J}{D} P_m^{non-full}\rho_w, \quad (37)$$

where the second term in the expression for $W_J^+$ represents divisions of wild-type cells that result in a mutation (probability $u$ per division). Solving the equation

$$W_j^+ = W_J^-,$$

we can find the value, $J$, that corresponds to selection-mutation balance:

$$J_{sel-mut} = \frac{Du}{\epsilon P_m^{non-full}(1-\mathcal{F})}, \quad \mathcal{F} = \frac{N_m P_w^{non-full} d_m \rho_m}{N_w P_m^{non-full} d_w \rho_w} \tag{38}$$

(note that the expression for mutant deme fitness, $\mathcal{F}$, is the same as appears in table 4 for this case).

Figure S7 illustrates this with an example. We ran stochastic simulations in a system of 100 demes with $K = 20$ each, in the presence of rare migrations and mutations. The mutants are assumed to have a negative selection coefficient ($s = -0.01$). Panel (a) of figure S7 shows a typical trajectory; the number of mutants in the system undergoes large fluctuations. Finding the temporal mean number of mutants of 516 independent simulations of this type, we plotted a histogram of mean mutant populations (panel (b)). The red vertical line shows the theoretical prediction, $J_{sel-mut}$, obtained by equation (38) (the red vertical line). We can see that the theory is consistent with the simulations, which is further illustrated in panel (c), where the mean temporal course of 516 simulations is plotted together with the predicted mean number of mutants.

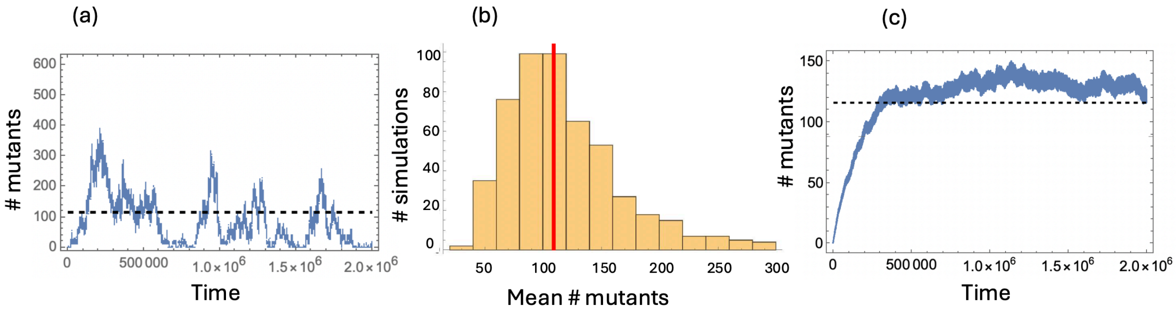

Figure S7: Selection-mutation balance. (a) A typical trajectory obtained by Gillespie simulations of a deme-structured population with mutations. The number of mutants is shown as a function of time. (b) A histogram of temporal averates of mutant numbers for 516 independent simulations. (c) Ensemble-averaged time-series for the number of mutants calculated from the 516 simulations, together with the standard error. The red vertical line and the dashed horizontal lines show the theoretical prediction, $J_{sel-mut}$, obtained from eqiation (38). The parameters are $D = 100, K = 20, s = -0.01, r_w = 5, d_w = 0.5, \epsilon = 10^{-4}, u = 10^{-6}, \tau = 1$.

#### 2.2 The reversal of the selection forces as a result of a changing turnover factor

A change in the turnover factor, $\tau$, can lead to qualitative changes in the behavior of a spatial or fragmented system, such that a disadvantageous mutant ($s < 0$) starts behaving as if it is selected for and takes over the population. This happens if the turnover factor is small enough. To see this, we observe that the expression $(1-\mathcal{F})$ appears in the denominator of equation (38). An increase of $\mathcal{F}$ (which results from a decrease in $\tau$) increases the level at which mutants are maintained in the system, until $\mathcal{F}$ grows above 1, at which point equation (38) breaks down and the mutant is no longer disadvantageous. This is illustrated in figure S8(a), where the singularity of the quantity $J_{sel-mut}$ is observed around $\tau \approx 0.2$.

Similarly, an advantageous mutant that takes over the population in a nearly-deterministic fashion with $\tau = 1$, may become disadvantageous if it is accelerated sufficiently (see panel (b) of figure S8).

The value of $\tau$ that reverses selection forces is given implicitly by the equation

$$\frac{\rho_m}{\rho_w} = \frac{N_w P_m^{non-full} d_w}{N_m P_w^{non-full} d_m}, \tag{39}$$

where the right hand side does not depend on $\tau$.

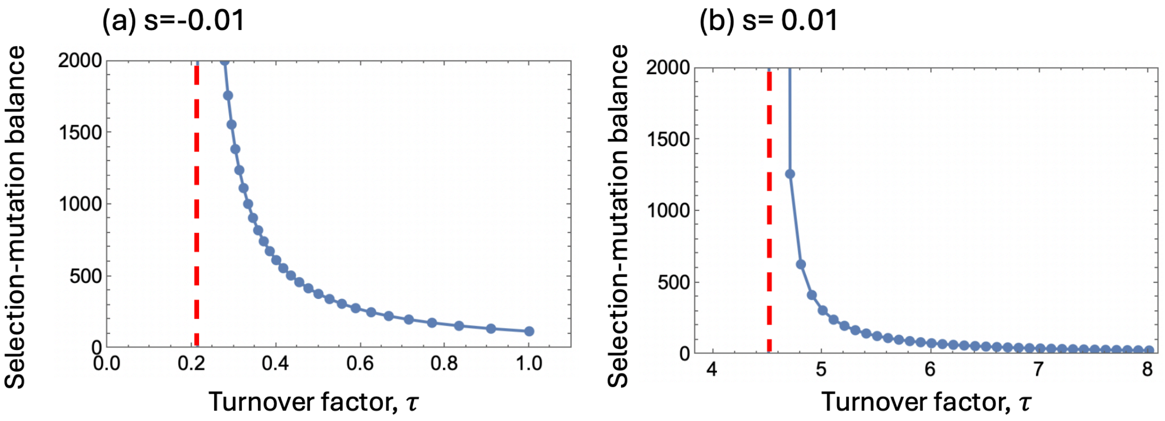

Figure S8: The effect of the turnover factor on mutant behavior in the presence of *de novo* mutations. The theoretically calculated selection-mutation balance ($J_{sel-mut} K(1 - d_m/r_m)$, equation (38), is plotted as a function of $\tau$. (a) Disadvantageous mutant, $s = -0.01$. (b) Advantageous mutant, $s = 0.01$. The red vertical line indicates the value of $\tau$ that makes the denominator in $J_{sel-mut}$ zero (equation (39). Other parameters are $K = 20, d_w/r_w = 0.1, \epsilon = 10^{-4}, u = 10^{-6}$.

### 3 Agent-based modeling

#### 3.1 Mutant fixation probability

Figure S9 illustrates the results of ABM simulations of a spatial system. It presents the dependence of the mutant fixation probability in a spatially explicit ABM on various system parameters. This figure should be compared with figure 2 of the main text, where we show similar results for the fragmented deme model.

#### 3.2 Dynamics in the presence of *de novo* mutations

Figure S10 shows the reversal of the selection force for mutants in spatially explicit ABM simulations. Panels (a,b) describe mutants with a negative selection coefficient ($s = -0.001$), which fluctuate around a selection-mutation balance if $\tau = 1$ (panel (a)), but take over the population if their division and death rates are slowed down compared to the wild-type by a factor $\tau = 0.1$ (panel (b)). In this case, they act as if they are selected for, although

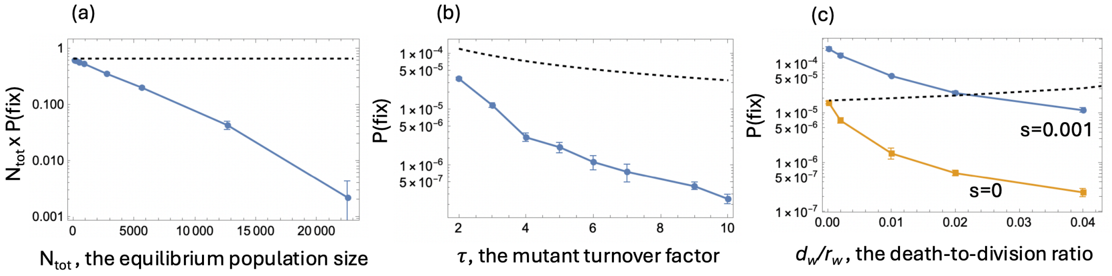

Figure S9: Spatial simulation results for the ABM model. In each plot, the mean and standard deviation is presented for each point. The dashed black line is the result for equivalent parameter values in a non-spatial system (with $s = 0$). (a) The dependence of $N_{tot}P(\text{fix})$ on the system size, $N_{tot}$, with $\tau = 2$. (b) The dependence of $P(\text{fix})$ on the turnover factor, $\tau$. The equilibrium population size is $N_{tot} = 5643$. (c) The dependence of $P(\text{fix})$ on the death-to-division ratio, $d_w/r_w$, for mutants with $\tau = 10$ and $s = 0.001$ (blue) and $s = 0$ (yellow). In all panels, $d_w/r_w = 0.04$ and $s = 0$ unless otherwise noted. The total number of independent runs is at least $10^6$ in panel (c) with $s = 0.001$ and at least $10^7$ in the rest of the cases.

the selection coefficient is negative and the life-time reproductive output of the mutants is smaller than that of the wild types.

Panels (c,d) show the opposite effect. There, the mutants are characterized by a positive selection coefficient ($s = 0.001$), and proceed to take over the population in a typical simulation corresponding to $\tau = 1$ (panel (c)). If however we inrease the turnover factor $\tau = 3$, we observe a qualitative change in the behavior, where mutants are maintained at a selection-mutation balance (panel (d)), despite them being characterized by a higher reproductive outcome compared to the wild-type individuals. Figure S10 should be compared to figure 3 of the main text, which shows similar effects for a deme-structured population.

### References

[1] Pierre-François Verhulst. Notice sur la loi que la population suit dans son accroissement. *Correspondence mathematique et physique*, 10:113–129, 1838.

[2] Raymond Pearl and Lowell J Reed. On the rate of growth of the population of the United States since 1790 and its mathematical representation. *Proceedings of the national academy of sciences*, 6(6):275–288, 1920.

[3] Ingemar Nåsell. Extinction and quasi-stationarity in the Verhulst logistic model. *Journal of Theoretical Biology*, 211(1):11–27, 2001.

[4] Charles R Doering, Khachik V Sargsyan, and Leonard M Sander. Extinction times for birth-death processes: Exact results, continuum asymptotics, and the failure of the Fokker–Planck approximation. *Multiscale Modeling & Simulation*, 3(2):283–299, 2005.

[5] Todd L Parsons and Christopher Quince. Fixation in haploid populations exhibiting density dependence II: The quasi-neutral case. *Theoretical population biology*, 72(4):468–479, 2007.

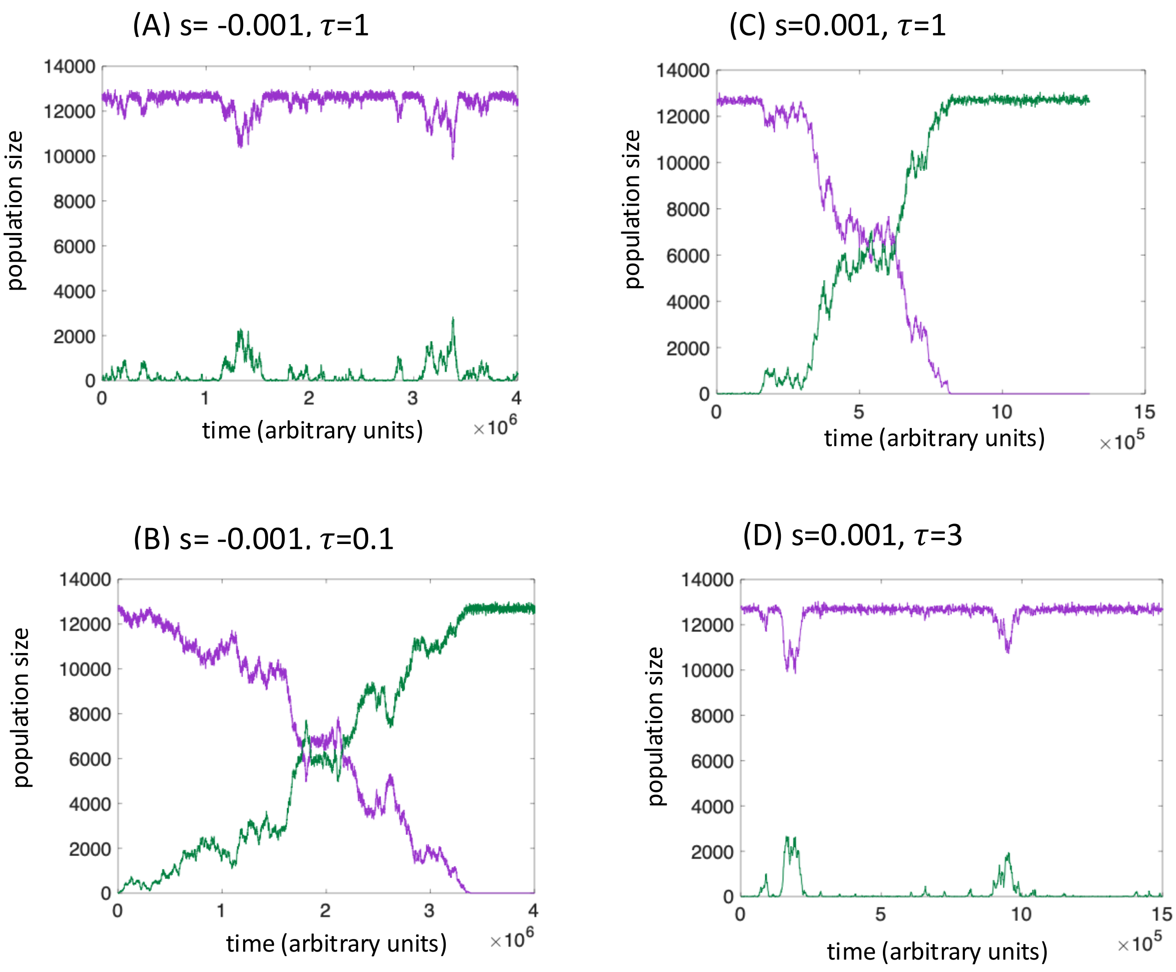

Figure S10: The reversal of selection forces in spatial ABM simulations. Time-series of the wild-type (purple) and mutant (green) numbers are shown for typical simulation realizations. (a) $s = -0.001, \tau = 1$ (selection-mutation balance); (b) $s = -0.001, \tau = 0.1$ (mutant takes over); (c) $s = 0.001, \tau = 1$ (mutant takes over); (d) $s = 0.001, \tau = 3$ (selection-mutation balance). Other parameters are $R_w = 0.05, D_w = 0.02, u = 10^{-5}$, and the grid size is $150 \times 150$.

[6] Todd L Parsons, Christopher Quince, and Joshua B Plotkin. Some consequences of demographic stochasticity in population genetics. *Genetics*, 185(4):1345–1354, 2010.

[7] Dominik Wodarz, Ajay Goel, and Natalia L Komarova. Effect of cell cycle duration on somatic evolutionary dynamics. *Evolutionary Applications*, 10(10):1121–1129, 2017.

[8] Dominik Wodarz and Natalia L Komarova. Mutant fixation in the presence of a natural enemy. *Nature Communications*, 14(1):6642, 2023.

[9] Ben Adlam, Krishnendu Chatterjee, and Martin A Nowak. Amplifiers of selection. *Proceedings of the Royal Society A: Mathematical, Physical and Engineering Sciences*, 471(2181):20150114, 2015.
